## Supplementary Information for "Non-destructive transcriptomics *via* vesicular export"


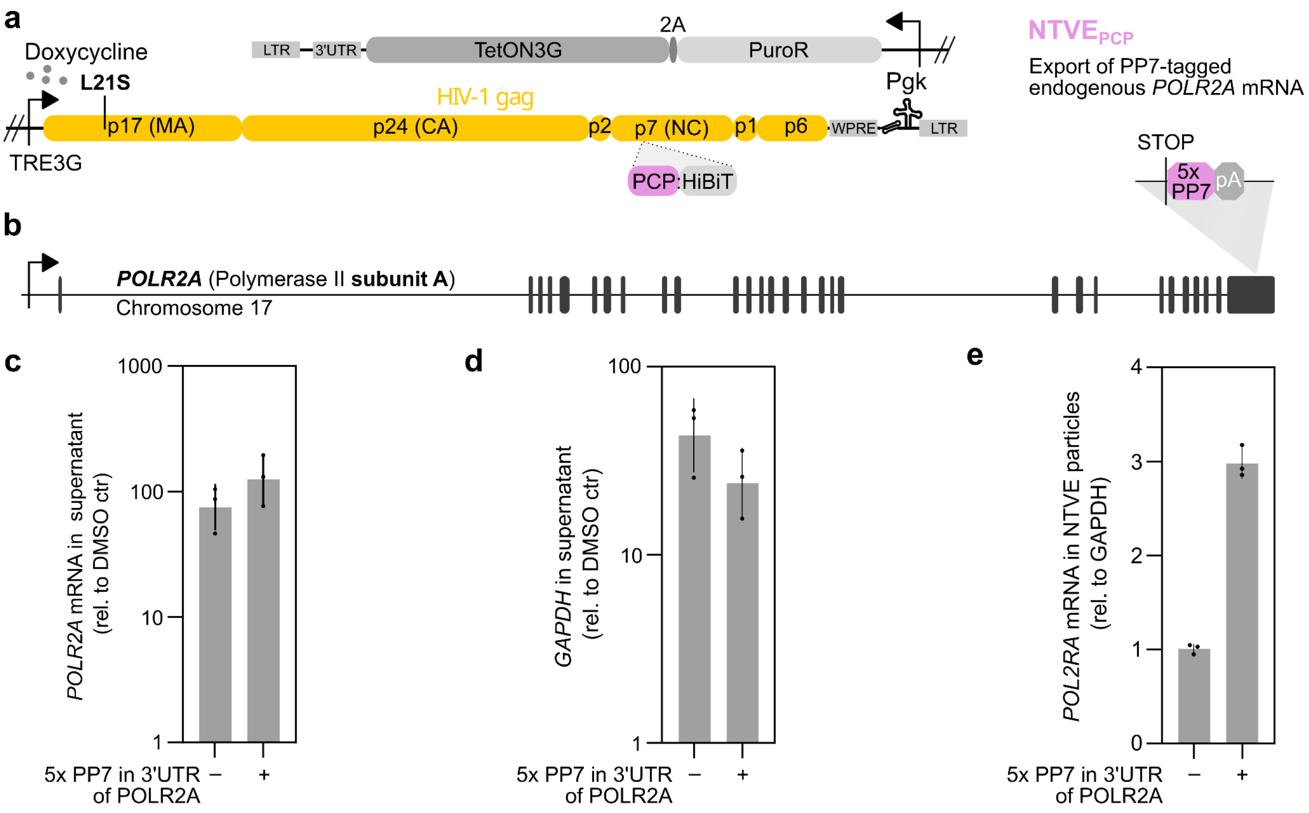


**Supplementary Figure 1: Tagging endogenous mRNA with PP7 for NTVE$_{PCP}$ export.**
**(a)** The PP7 coat protein (PCP, ~0.4 kbp) serves as a specific adapter to export PP7-tagged barcode RNAs. **(b)** *POLR2A* was chosen as an exemplary locus for integration of five concatenated PP7 aptamers *via* CRISPR/Cas9, positioned in the 3′UTR of the transcribed mRNA. **(c)** RT-qPCR quantification of NTVE-mediated export of *POLR2A* transcripts from an NTVE$_{PCP}$ cell line with a PP7-tagged *POLR2A* 3′UTR compared with a standard NTVE$_{PCP}$ cell line. Cells were induced with 500 ng/ml dox (*vs.* DMSO control). **(d)** Non-specific RNA export assessed by endogenous *GAPDH* abundance in the NTVE$_{PCP}$ fraction of both cell lines. **(e)** *POLR2A*-to-*GAPDH* ratio in both NTVE fractions after dox induction. Bars represent the mean of three biological replicates, with error bars indicating the *s.d.* Source data are provided as a Source Data file.

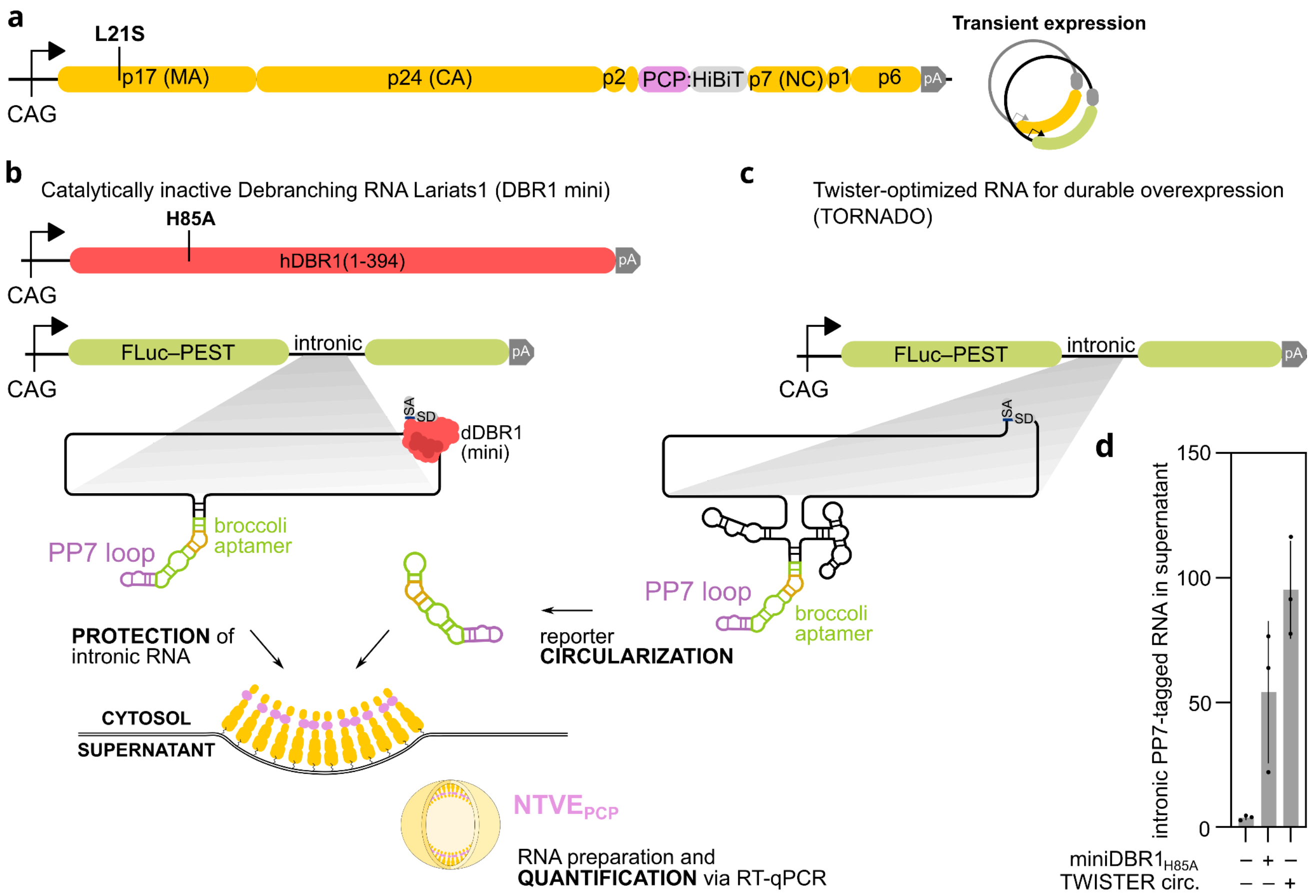


**Supplementary Figure 2: Encoding of RNA barcodes in introns of Polymerase II-transcribed constructs for cellular export *via* NTVE$_{PCP}$.**

**(a)** Genetic construct for transient expression of NTVE$_{PCP}$ harboring a HiBiT tag for bioluminescent quantification of export efficiency. **(b)** Construct encoding a catalytically inactive (H85A) and truncated debranching enzyme (miniDBR1$^{H85A}$) to protect an intronic sequence encoding a PP7-tagged Broccoli aptamer from degradation. **(c)** Intron-encoded PP7-tagged Broccoli aptamer that undergoes circularization *via* the Tornado system. **(d)** To promote the accumulation of otherwise unstable intronic RNA and enable its export, the PP7-tagged intronic RNA was stabilized by either co-expression of catalytically inactive DBR1 or by the Tornado system. RNA levels were quantified by RT-qPCR and normalized to a control condition lacking NTVE$_{PCP}$ expression.  Source data are provided as a Source Data file.

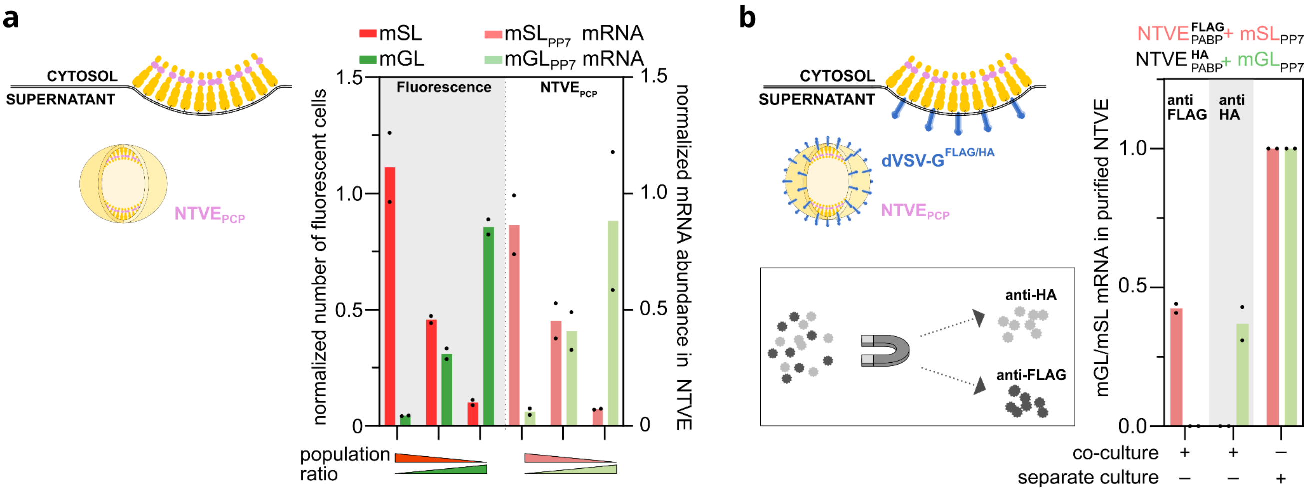


**Supplementary Figure 3: Relative reporter abundance in supernatant of NTVE$_{PCP}$ co-cultures and epitope-specific purification of NTVE$_{PCP}$ particles.**

**(a)** HEK293T cells with an integrated NTVE$_{PCP}$ expression cassette were transfected with plasmids encoding either mScarlet-I (mSL) or mGreenLantern (mGL) with PP7-tagged 3′UTRs. Twenty-four hours post-transfection, cells were reseeded at ratios of 10:1, 1:1, and 1:10. Co-cultured populations were subsequently quantified by fluorescence microscopy (left *y*-axis). In addition, mGL and mSL mRNA abundance in the supernatant was quantified by RT-qPCR (right *y*-axis). Both measures were normalized to monocultures expressing only one of the reporters ($n$ = 2). **(b)** For purification using epitope-tagged dVSV-G(W72A), stable NTVE$_{PCP}$ HEK293T cells were transfected with plasmids encoding either mSL or mGL with PP7-tagged 3′UTRs. Twenty-four hours post-transfection, cells were reseeded at a 1:1 ratio, followed by NTVE induction (500 ng/ml dox). Tagged VLPs from the co-cultured cell lines were enriched with either anti-FLAG or anti-HA magnetic beads, and mGL/mSL abundance was quantified by RT-qPCR ($n$ = 2). Source data are provided.

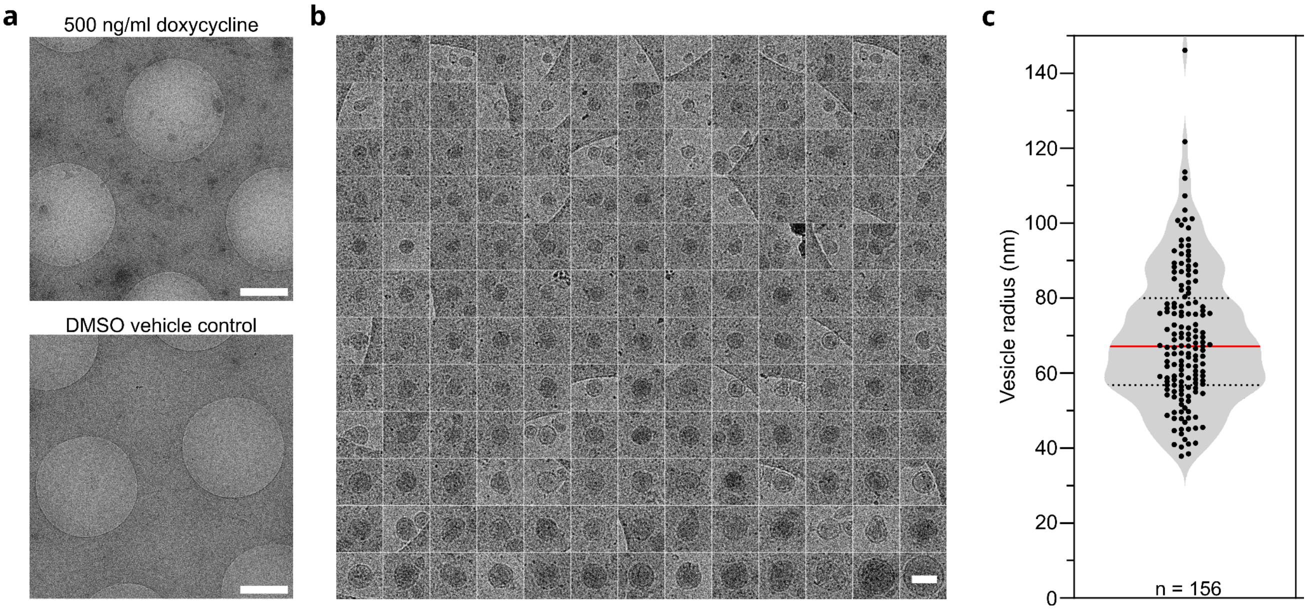


**Supplementary Figure 4: Cryo-TEM of exported NTVE$_{PABP}$ vesicles.**

**(a)** Low-magnification cryo-TEM overviews showing relative particle abundances after dox induction (top) and DMSO vehicle control (bottom). Scale bars: 1 µm. **(b)** Montage of NTVE$_{PABP}$ vesicles, purified *via* ultracentrifugation from the supernatant of HEK293T cells 72 h post-dox induction. The scale bar indicates 200 nm. **(c)** Distribution of estimated vesicle radii from **(b)**. Radii were computed as those of circles with an equivalent area equal to that of the manually segmented vesicles, to account for deviations from circularity in some particles. The red line indicates the median (67 nm, $n$ = 156). Comparable distributions are obtained by fitting circles to the vesicle boundaries or measuring the longest-axis diameter. Source data are available.

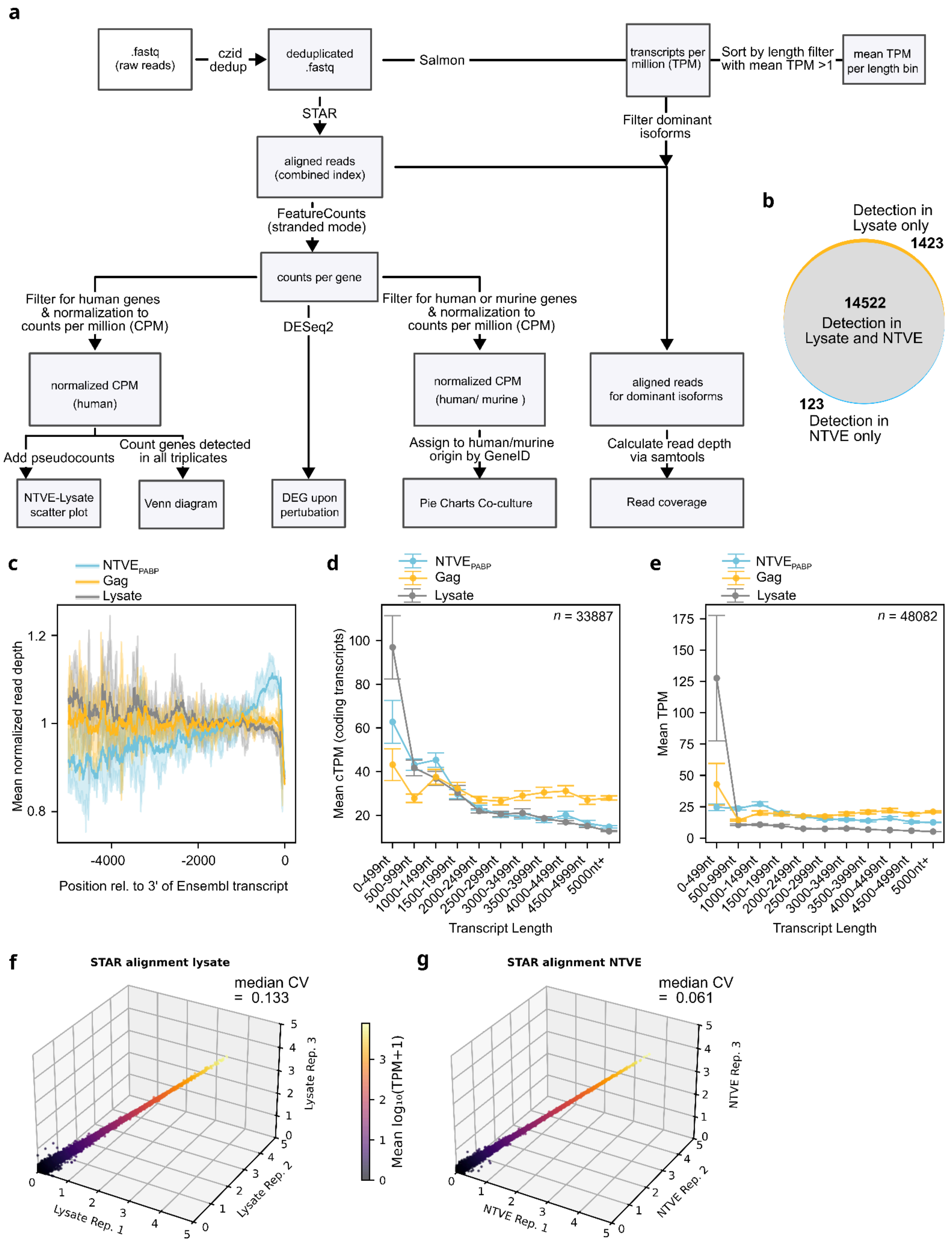


**Supplementary Figure 5: Additional NTVE characterization.**
**(a)** Alignment pipeline for paired-end RNA-seq. Reads were aligned *via* STAR (Spliced Transcripts Alignment to a Reference, v2.7.10b_alpha_220111) to a combined human–murine index. Uniquely mapped fragments were used to calculate read counts with featureCounts (v2.0.3). After quantification, reads were filtered for human or murine origin and normalized to counts per million (CPM).

Pseudocounts of $10^{-3}$ were added to ensure numerical stability. Differential gene expression was analyzed with the DESeq2 package (v1.34.0) in R (v4.1.2). Salmon (v1.10.1) was used to compute transcripts per million (TPM) for each transcript and to plot the mean TPM per length bin. For read coverage analysis, STAR-aligned reads were filtered for dominant isoforms identified by Salmon (*i.e.*, >90% of total TPM of the associated gene) and aligned to the 3′ end of the Ensembl-derived transcript annotation. Read depth was calculated with SAMtools (v1.19.2) using the `depth` command for regions corresponding to the sample-specific set of dominant isoforms. **(b)** Venn diagram showing the total count of all genes with at least one read in all three replicates in the lysate or the NTVE fraction. A list of genes detected in the lysate but not in NTVE is given in **Supplementary Information 2**. **(c)** Average normalized read depth from $NTVE_{PABP}$ (blue) and Gag without an adapter (yellow), plotted as a function of the nucleotide position relative to the 3′ poly(A) site. As a reference (gray), a lysate was subjected to subcellular fractionation to isolate the cytoplasmic fraction and remove nuclear components. Lines represent the mean ± 95% CI of triplicates. **(d)** Mean coding transcripts per million (cTPM) of expressed (TPM > 1) transcripts, binned by transcript length. Lines represent the mean ± *s.d.* of triplicates. **(e)** Mean transcripts per million (TPM) of expressed (TPM > 1) transcripts, binned by transcript length. Lines represent the mean ± *s.d.* of triplicates. **(f)** Correlation plots of $\log_{10}$-transformed CPMs of NTVE (triplicates) exported over 72 hours and **(g)** transcripts from the corresponding lysate obtained at the endpoint. Coefficients of variation (CV) are indicated above each plot. Samples for **(b–d)** were prepared without enrichment (total RNA-seq), followed by rRNA depletion, while samples in **(e–g)** were poly(A)-enriched before reverse transcription. Raw sequencing data and code are available *via* Zenodo.

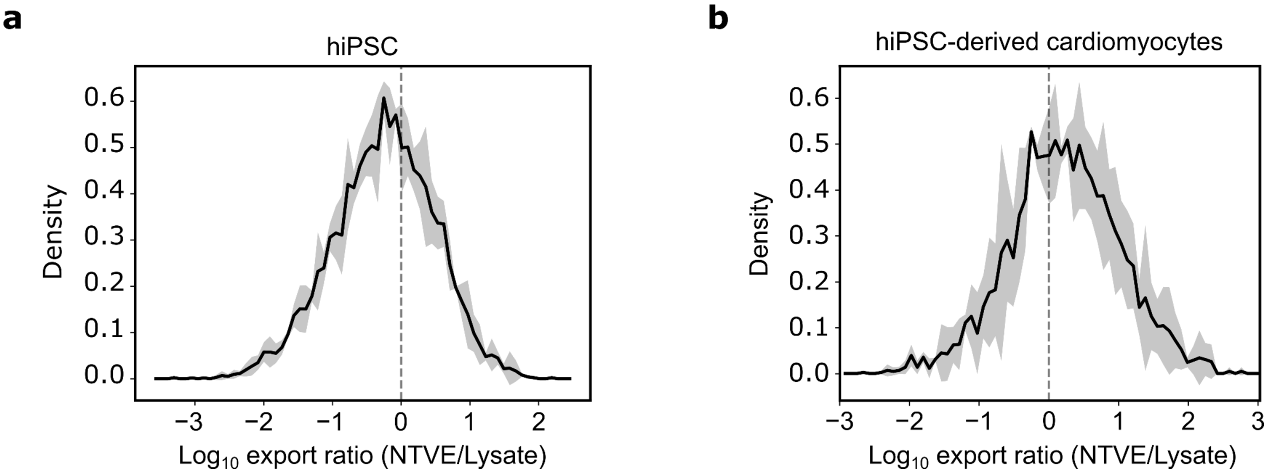


**Supplementary Figure 6: Distribution of NTVE export ratios.**
**(a,b)** Distribution of the export ratio across all endogenous protein-coding genes in **(a)** MRIi003-A hiPSCs and **(b)** MRIi003-A hiPSC-derived cardiomyocytes. The line represents the mean density of three replicates, with the gray shading indicating the 95% confidence interval. Raw sequencing data and code are provided *via* Zenodo

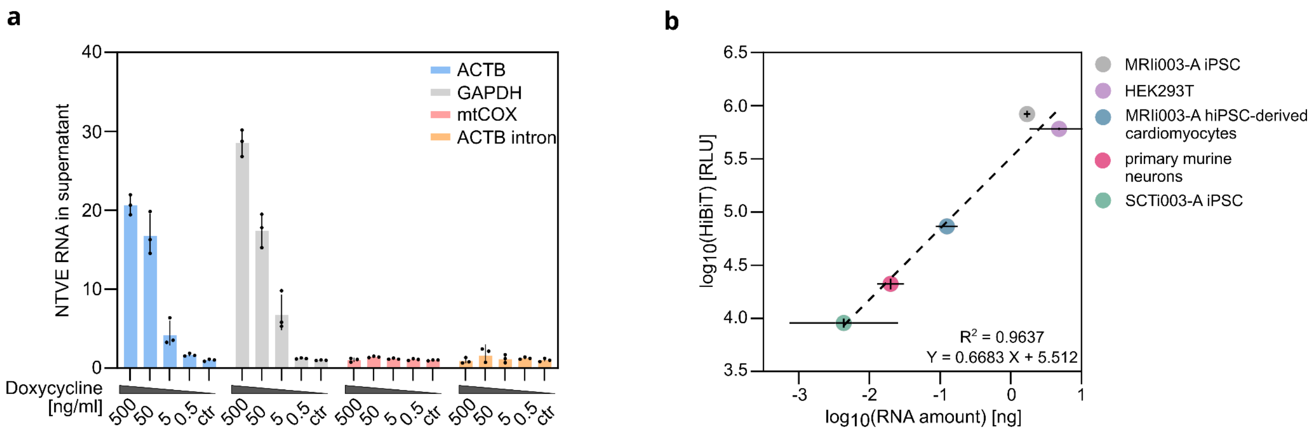


**Supplementary Figure 7: Optimization of dox-induction of $NTVE_{PABP}$ in different cell lines.**
**(a)** Dynamic range of dox-induced $NTVE_{PABP}$ export of two housekeeping transcripts (*ACTB* and *GAPDH*) measured by RT-qPCR 72 hours after induction. As negative controls, we selected two polyadenylated transcripts not expected to be accessible for cytosolic export: *mt-CO1*, which is encoded on the mitochondrial genome, and an intron of *ACTB*. The TetON3G transactivator was driven by a PGK promoter. **(b)** Correlation between the bioluminescent signal from the HiBiT reporter tag fused to $NTVE_{PABP}$ and the RNA amount in the supernatant 72 hours after induction. Data were obtained from several cell lines, seeded at different densities and plate formats. In **(a)** and **(b)**, three biologically independent replicates are presented as mean ± *s.d.* Source data are provided.

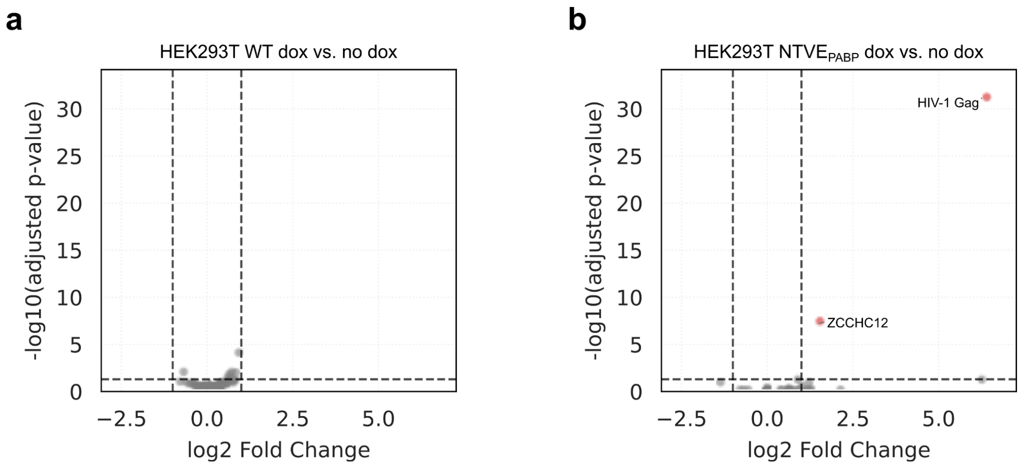


**Supplementary Figure 8: Dox effects on differential gene expression in lysates.**
**(a)** Volcano plot of differentially expressed genes in wild-type HEK293T cells after induction with 500 ng/ml dox. **(b)** Volcano plot of differentially expressed genes in the HEK293T $NTVE_{PABP}$ cell line after dox induction compared with DMSO vehicle control (ecotropic lentiviral delivery; Pgk promoter-driven TetON3G). Raw sequencing data and code are deposited at Zenodo.

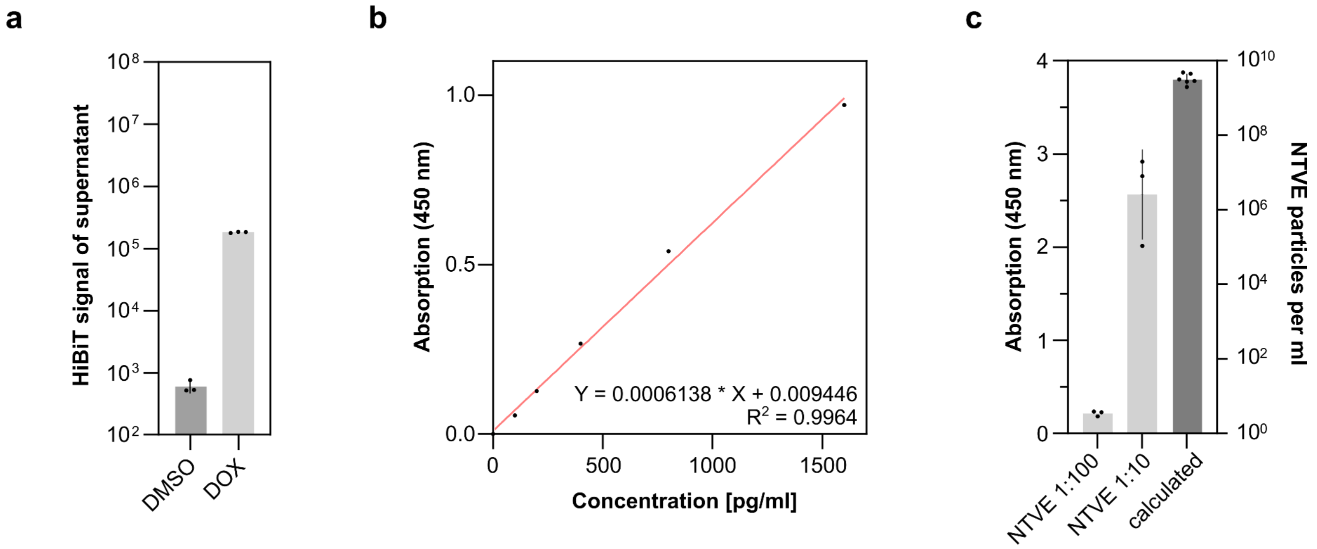


**Supplementary Figure 9: Quantification of NTVE vesicle titers 72 hours post dox induction.**
**(a)** HiBiT-mediated luminescence signal from the supernatant of the dox-induced (500 ng/ml) NTVE HEK293T cell line compared with a DMSO control condition. **(b)** Quantification of NTVE particle titers by anti-p24 (HIV-1) ELISA with a p24 standard. **(c)** ELISA measurements of two dilutions of supernatants (*n* = 3) and calculation of the corresponding particle numbers, assuming 2,500 monomers per particle. Source data are provided.

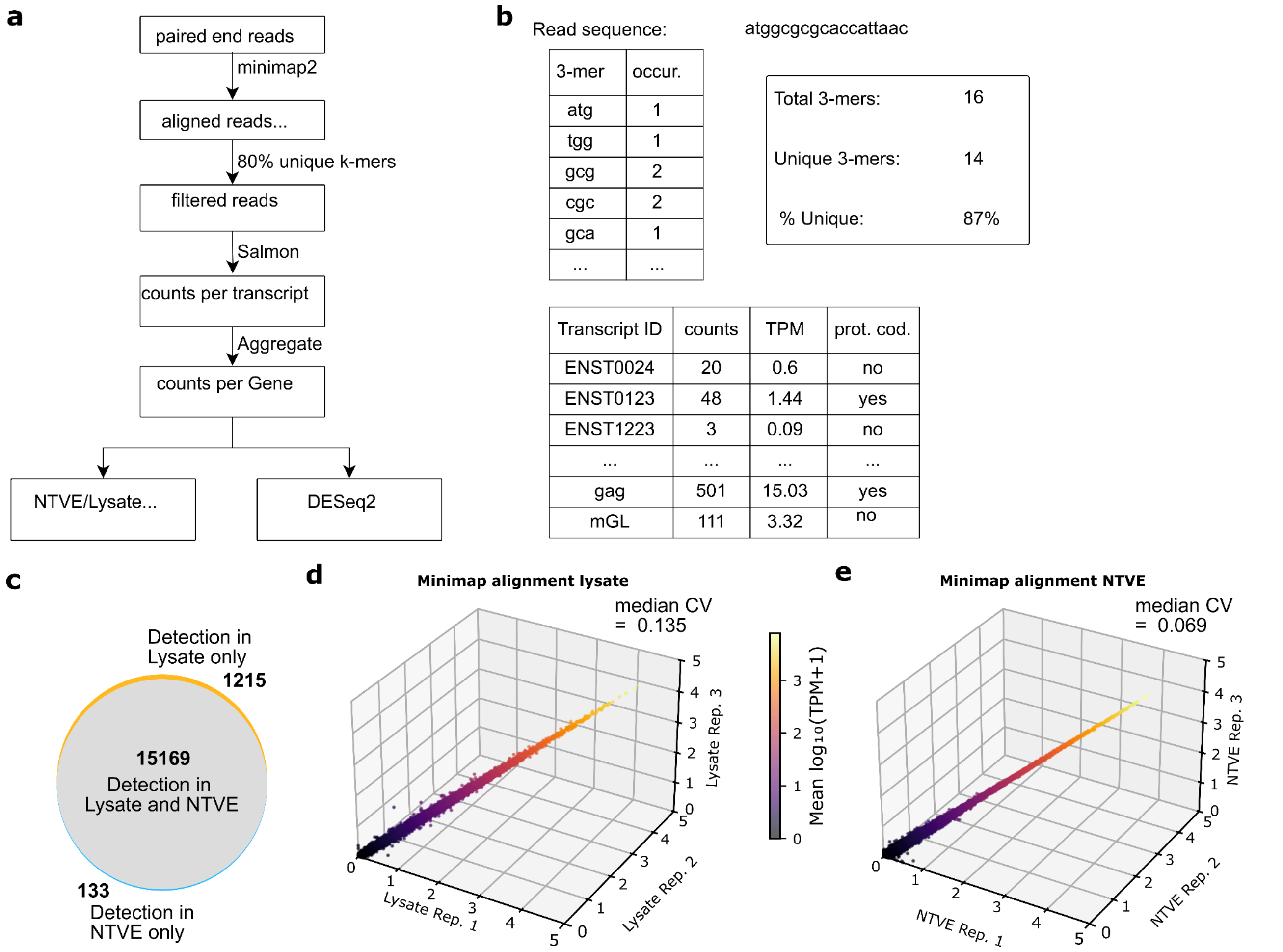


**Supplementary Figure 10: Computational pipeline and quality metrics for NTVE RNA-seq analysis.**
**(a)** Two-stage alignment and quantification pipeline for paired-end short-read RNA-seq. Minimap2 was used to align reads against the reference transcriptome. The unique k-mer cutoff was based on computing the proportion of uniquely occurring k-mers across all k-mers in each read. If this proportion fell below 80%, the read pair was excluded from further analysis. Salmon was used to quantify reads mapping to each transcript and normalize counts to transcripts per million (TPM). For downstream processing, only protein-coding transcripts were considered. **(b)** Schematic of the Salmon quantification output with appended transcript annotation. **(c)** Venn diagram of genes detected in lysate, NTVE, or both. **(d)** Correlation plot of $log_{10}$-transformed CPMs across NTVE triplicates exported over 72 hours. **(e)** Correlation plot of $log_{10}$-transformed CPMs across lysate triplicates. Coefficients of variation (CV) are indicated above each plot. Raw sequencing data and code are provided *via* Zenodo.

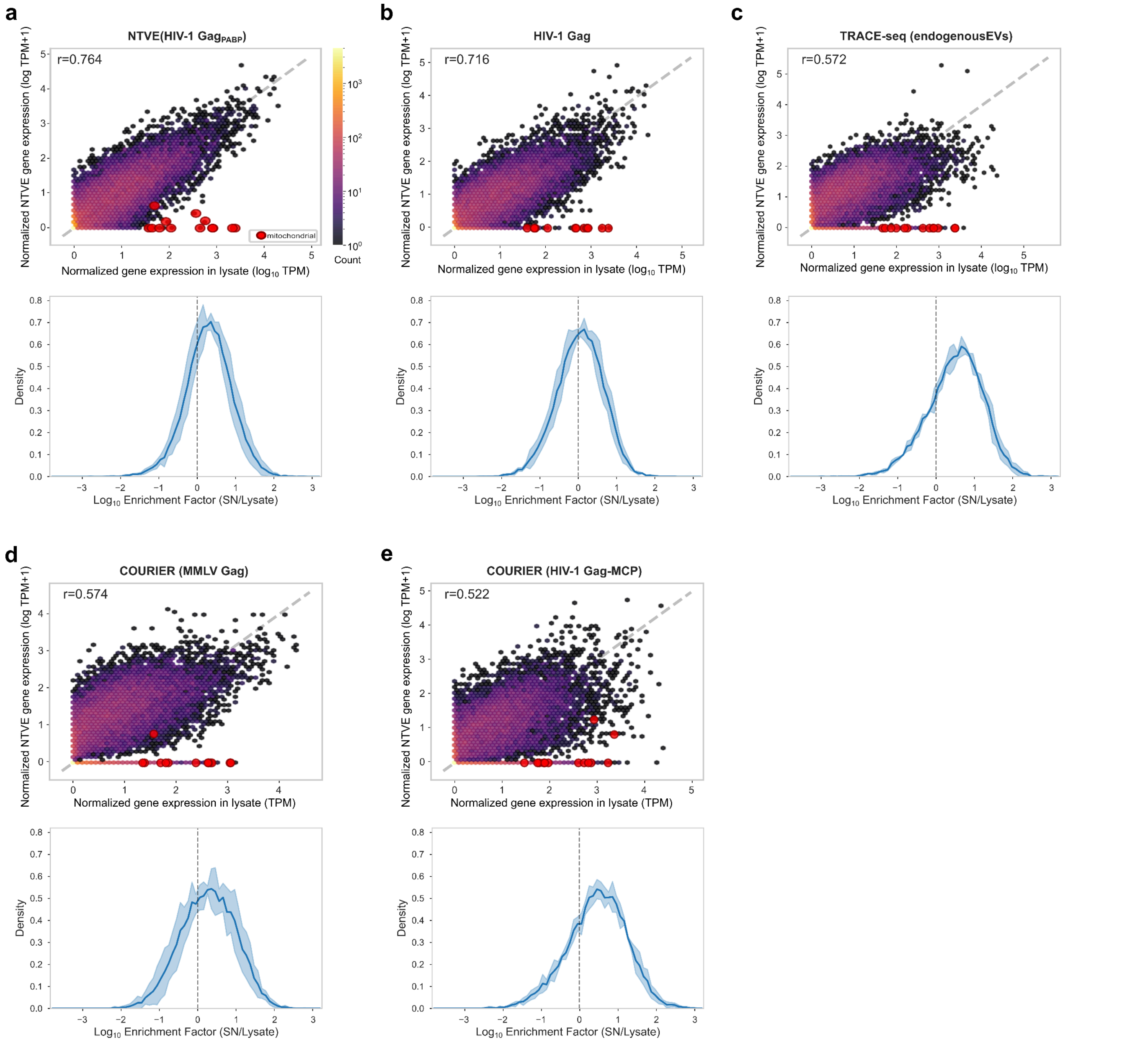


**Supplementary Figure 11: Transient transfection of HEK293T cells with different constructs to mediate RNA export.**
**(a–e)** Scatter plots of protein-coding gene-derived TPMs for supernatant and corresponding cell lysates three days post plasmid transfection, showing expression from different budding modules (top; $n$ = 1) and the corresponding export ratio (bottom; $n$ = 3). All export systems were expressed from a moderate-strength PGK promoter. RNA was isolated from the supernatant of cells expressing HIV-1 Gag$_{PABP}$ **(a)**, HIV-1 Gag **(b)**, endogenous EVs **(c)**, MMLV Gag **(d)**, and HIV-1 Gag-MCP **(e)** 72 hours post-transfection. Mitochondrial genes are indicated in red. Bottom: distribution of the export ratio across all endogenous protein-coding genes. The line represents the mean density, with the blue shading indicating the 95% confidence interval. Raw sequencing data and code available at Zenodo.

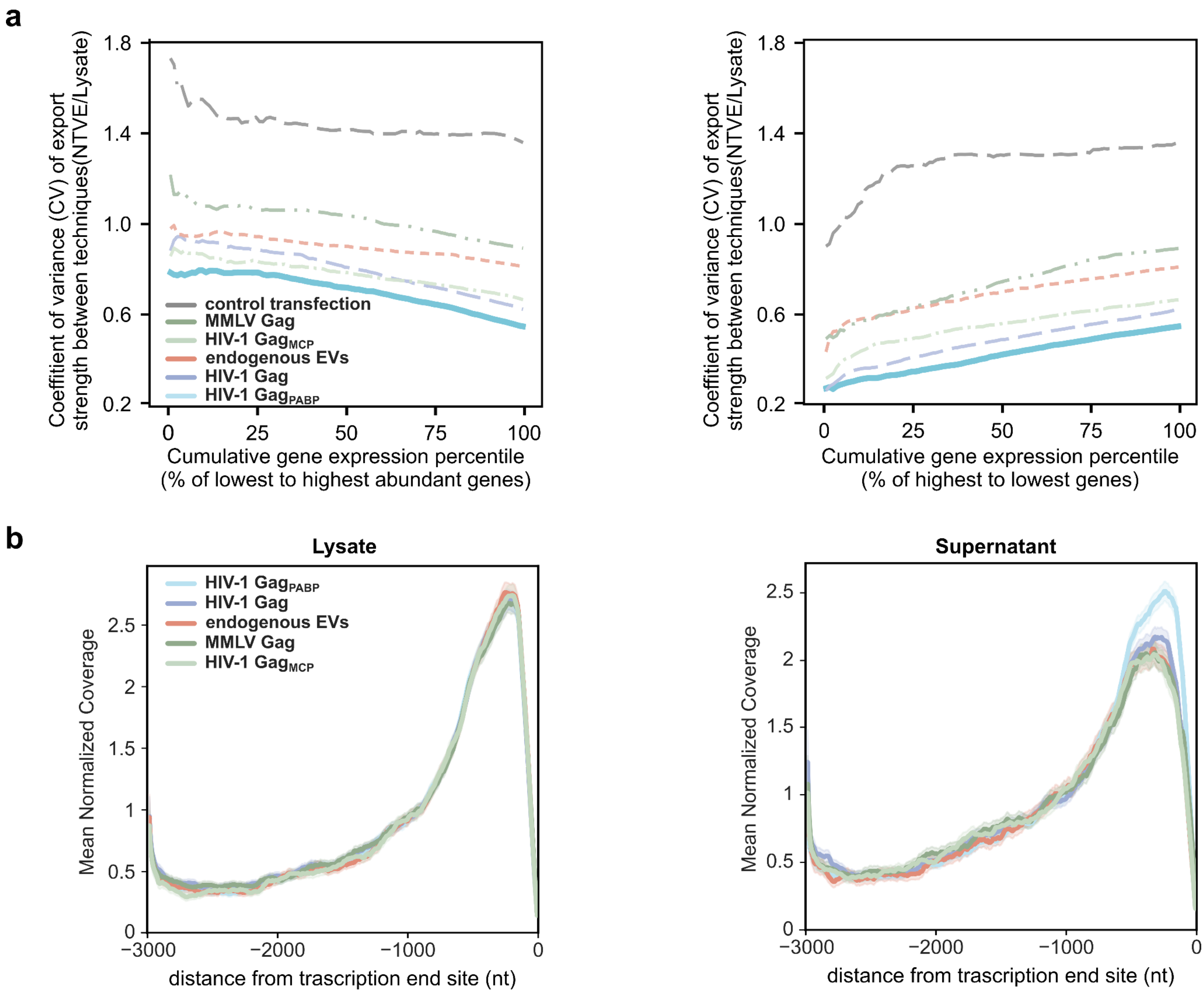


**Supplementary Figure 12: Reproducibility and read coverage of RNA export methods.**
**(a)** Coefficient of variation (CV) for different export methods plotted as a function of the cumulative percentile of detected genes, ranked from lowest to highest expression (left) or highest to lowest (right). The gray line represents a transfection control. **(b)** Average normalized read depth from $NTVE_{PABP}$ and other RNA export methods, plotted as a function of the nucleotide position relative to the 3′ poly(A) site. Lines represent the mean ± 95% confidence interval of triplicates. Raw sequencing data and code are provided *via* Zenodo.

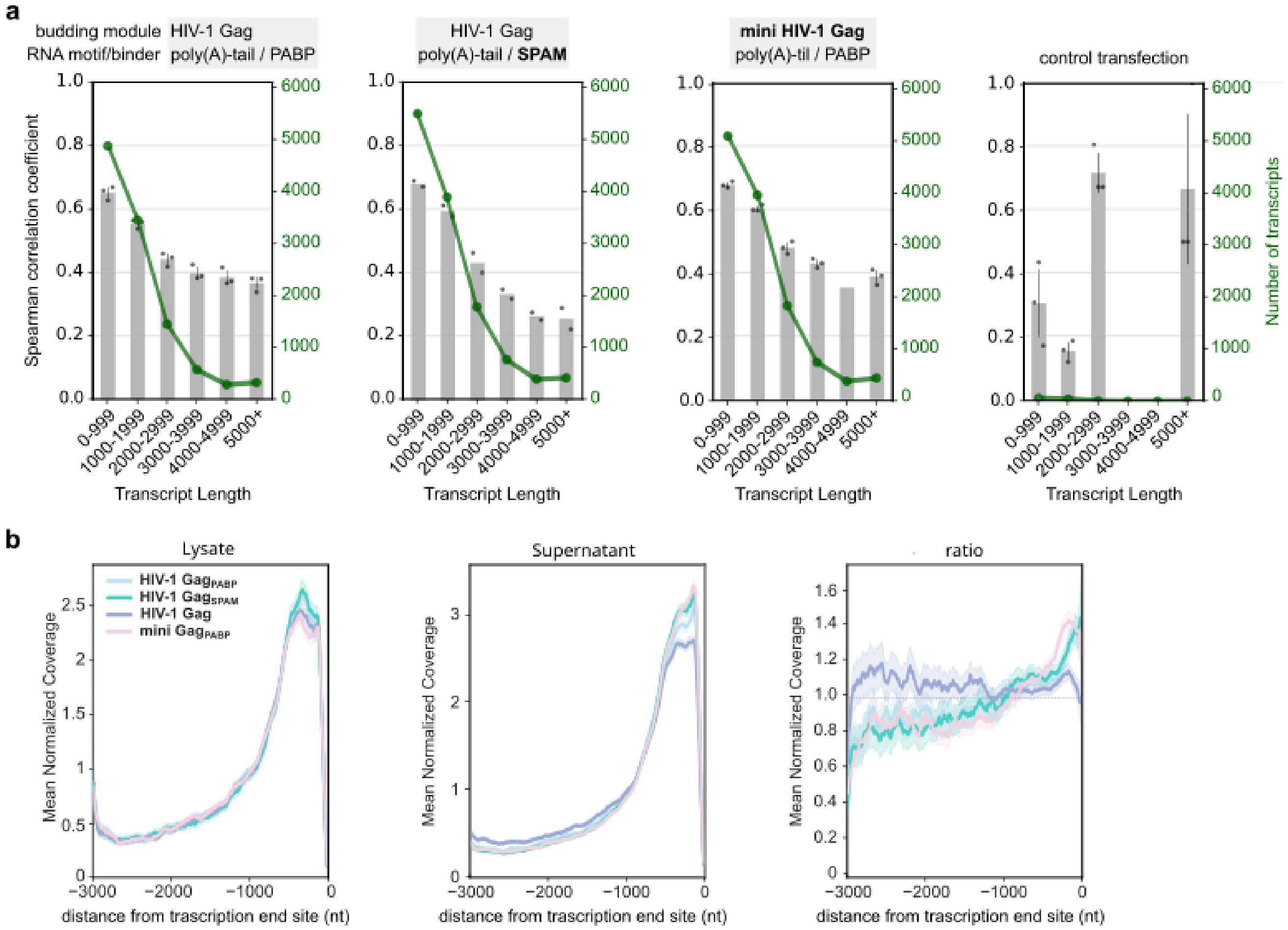


**Supplementary Figure 13: Engineered sPAM2 peptide as an alternative adapter for PABP-mediated mRNA recruitment.**
(**a**) Spearman correlation coefficients between supernatant and lysate transcript abundances, binned by transcript length, for HIV-1 $Gag_{PABP}$, HIV-1 $Gag_{sPAM2}$, and mini HIV-1 $Gag_{sPAM2}$. Grey bars represent mean coefficients (*n* = 3, n=2 for HIV-1 Gag sPAM2) with 95% confidence intervals; the green line indicates the mean number of detected transcripts per bin. The large confidence intervals in the control transfection with NanoLuc luciferase (no VLP formation) are due to the low number of detected transcripts. (**b**) Coverage profiles for lysate, supernatant, and the respective ratio. Coverage was computed from the 5,000 most abundant transcripts in each sample. Profiles were binned at 10 bp resolution from −3,000 bp upstream to the transcription end site (position 0). Lines represent mean normalized coverage across all transcript–replicate combinations, with shaded regions indicating 95% confidence intervals. Biological replicates within each sample were aggregated for statistical analysis. Raw sequencing data and code provided *via* Zenodo.

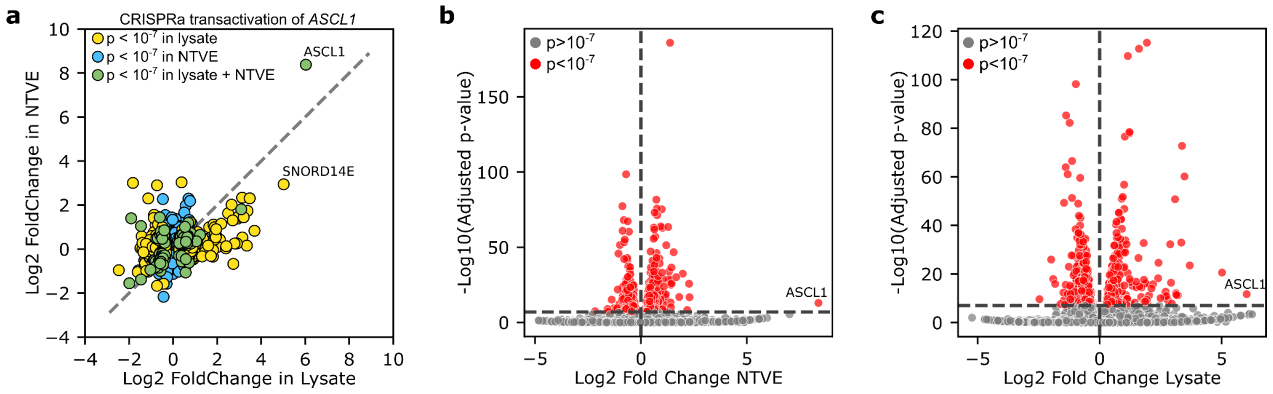


**Supplementary Figure 14: CRISPRa-driven transactivation of *ASCL1.***
**(a)** Correlation plot of significant fold changes between $NTVE_{PABP}$ and the corresponding lysate after export for 72 hours from cells with CRISPRa transactivation of *ASCL1* compared to cells without. Corresponding volcano plots for **(b)** $NTVE_{PABP}$ and **(c)** the corresponding lysate. Raw sequencing data and code are provided *via* Zenodo.

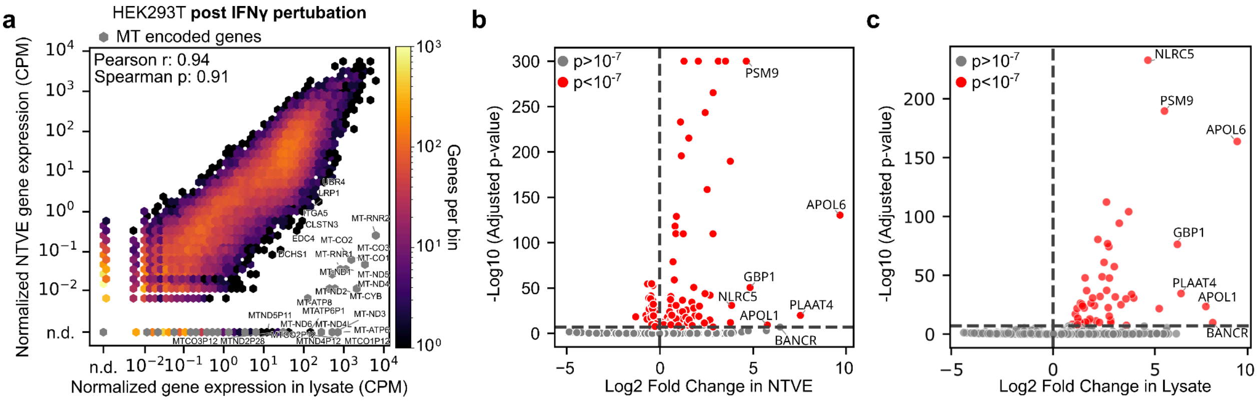


**Supplementary Figure 15: Additional characterization of the Interferon-γ perturbation experiment.**
**(a)** Normalized counts per million (CPM) for NTVE vesicles *vs.* corresponding lysates after dox induction and perturbation with IFN-γ. Genes encoded on the mitochondrial genome are labeled with their gene symbol (gray). Corresponding volcano plots for **(b)** $NTVE_{PABP}$ and **(c)** the corresponding lysate. Raw sequencing data and code are provided *via* Zenodo.

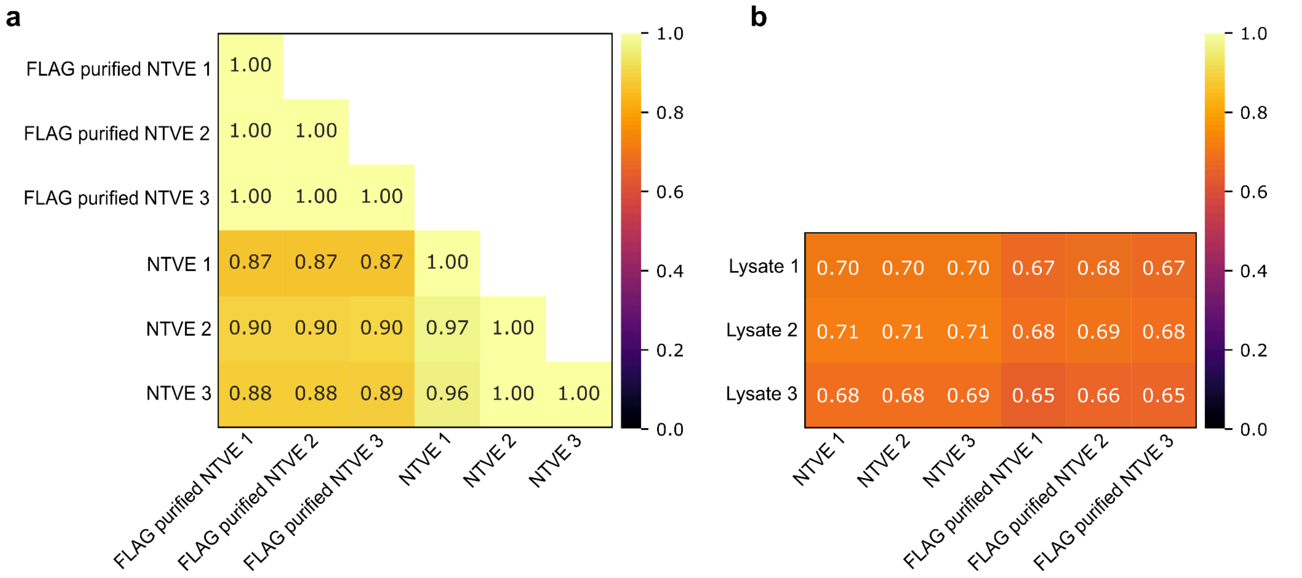


**Supplementary Figure 16: FLAG purification of NTVE particles from HEK293T cells.**
**(a)** Pearson correlations within replicates and across conditions for unpurified NTVE preparations and after FLAG purification. **(b)** Pearson correlation of the transcriptome obtained from whole-cell lysate with unpurified NTVE and FLAG-purified NTVE. Raw sequencing data and code available at Zenodo.

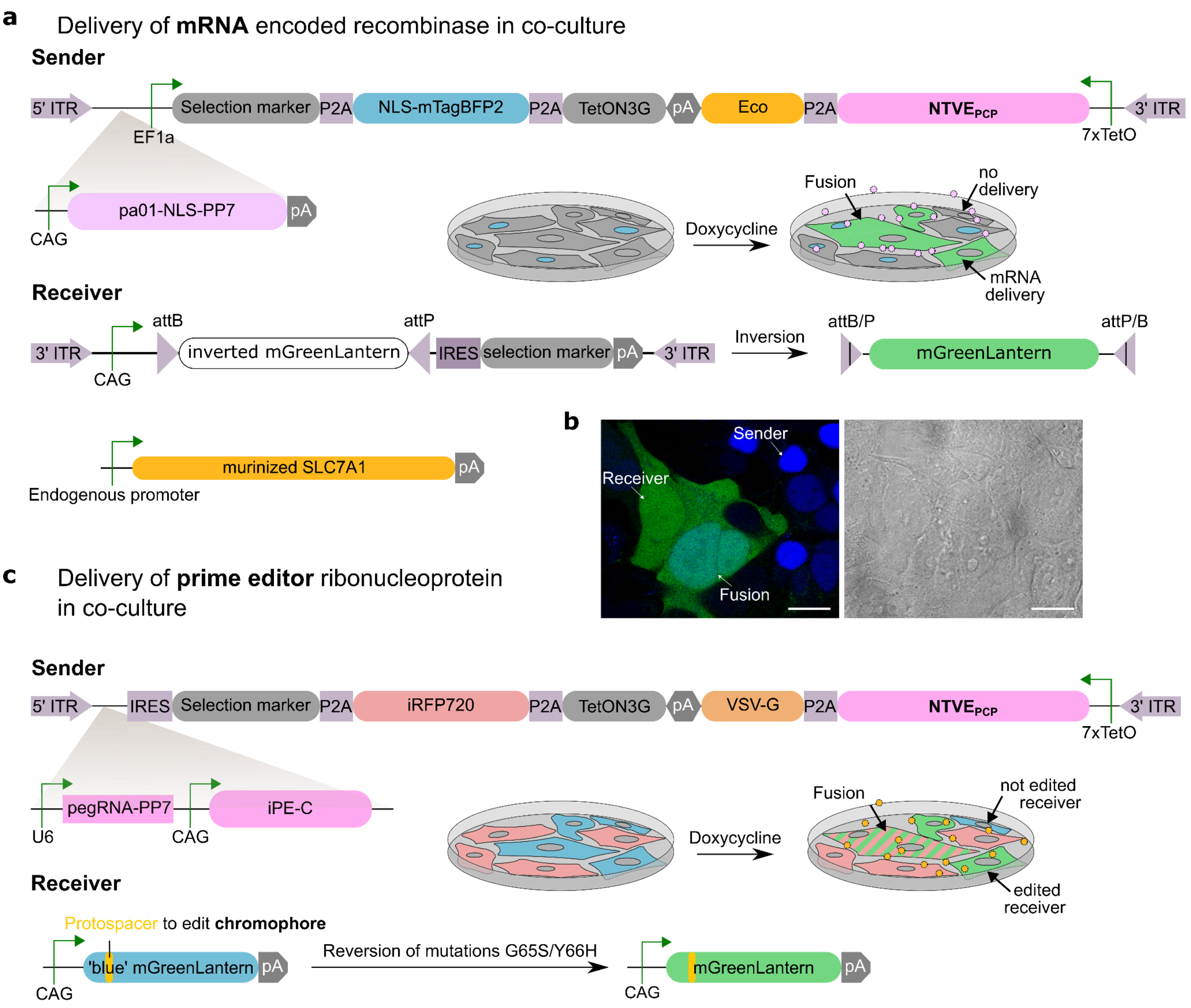


**Supplementary Figure 17: Genetic constructs for NTVE-mediated intercellular communication.**
**(a)** Schematic of the genetic construct for the cell-cell communication system in HEK293T cells based on an mRNA-encoded recombinase. The sender cells co-express three genetic elements: a PP7-tagged mRNA encoding the large serine recombinase (LSR) pa01, a nuclear-localized blue fluorescent protein, and dox-inducible NTVE$_{PCP}$ together with (*via* P2A) the Eco-ΔC16 glycoprotein for pseudotyping. The recipient cell expresses the murinized Eco-specific receptor SLC7A1 and carries the recombinase-responsive attB/attP cassette, which is inverted to express mGreenLantern upon pa01 activity. Both the sender construct and the reporter construct of the recipient cell are flanked by PiggyBac ITRs for genomic integration *via* PiggyBac transposase.
**(b)** Fluorescence microscopy showing blue sender and recipient cell nuclei with mGreenLantern expression. A fusion event (corresponding to the upper right quadrant in Figure 5c) is also visible, likely mediated by direct interaction of Eco and SLC7A1 on the respective cell surfaces.
**(c)** Schematic of the genetic construct for prime editor–based cell-cell communication in HEK293T cells. Here, a prime editor (iPE-C) is packaged together with a PP7-tagged pegRNA as a ribonucleoprotein (RNP). The recipient cell carries a reporter construct encoding a mGreenLantern mutant that exhibits blue fluorescence. The reporter contains a protospacer sequence that enables pegRNA-directed prime editing to revert the G65S/Y66H mutations, switching fluorescence from blue to green. In this configuration, sender cells express the far-red fluorescent protein miRFP670nano3 as a marker, and vesicles are pseudotyped with VSV-G glycoprotein.

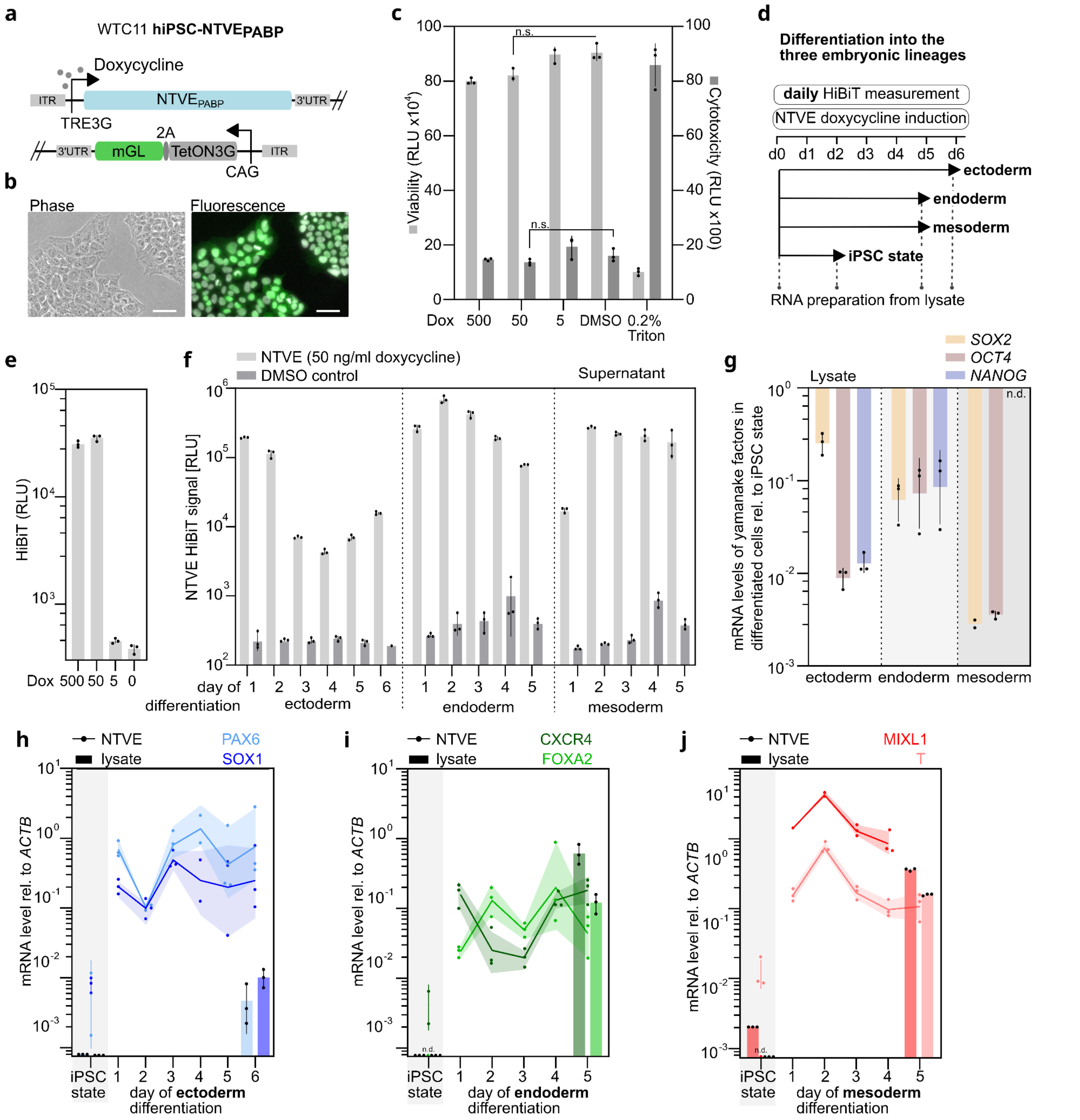


**Supplementary Figure 18: NTVE export levels during WTC11 hiPSC differentiation.**

**(a)** Schematic of the PiggyBac insertion cassette for human induced pluripotent stem cells (hiPSC) encoding a CAG-promoter-driven TetON3G to mediate doxycycline-dependent expression of $NTVE_{PABP}$ *via* a TRE3G promoter. The expression of TetON3G is coupled to that of mGreenLantern *via* a 2A peptide.

**(b)** Phase and fluorescence microscopy of WTC11 $NTVE_{PABP}$. Scale bar for microscopy images: 50µm.

**(c)** Viability and cytotoxicity assay of hiPSC at given doxycycline concentrations 48 hours post-induction.

**(d)** Timeline of the trilineage differentiation. NTVE-RNA was prepped daily while NTVE budding was maintained with 50 ng/mL doxycycline during differentiation to endo- and mesoderm (5 media exchanges) and ectoderm (6 media exchanges). RNA from lysates was prepped in the hiPSC state and as an endpoint measurement from the respective germ layer.

**(e)** Quantification of $NTVE_{PABP}$ budding efficacy at given doxycycline levels by HiBiT measurement from the supernatant 48 hours post-induction.

**(f)** HiBiT bioluminescence signals measured from the supernatant of hiPSCs (dox-induced *vs.* DMSO vehicle control) at the indicated days of differentiation into ectoderm, endoderm, and mesoderm, respectively.

**(g)** Relative mRNA expression of pluripotency markers after differentiation into the three germ layers compared to the undifferentiated hiPSC state. Bars represent the mean of three biological replicates; error bars indicate *s.d.*

**(h–j)** *ACTB*-normalized mRNA levels of lineage-specific markers in the supernatant over the time course of hiPSC differentiation towards ectoderm **(h)**, endoderm **(i)**, or mesoderm **(j)**. The terminal time point of *MIXL1* was excluded owing to an aberrant melting curve. In (c), (e–j), bars represent the mean of three biological replicates; error bars or shading indicate the *s.d.* Source data are provided as a Source Data file.

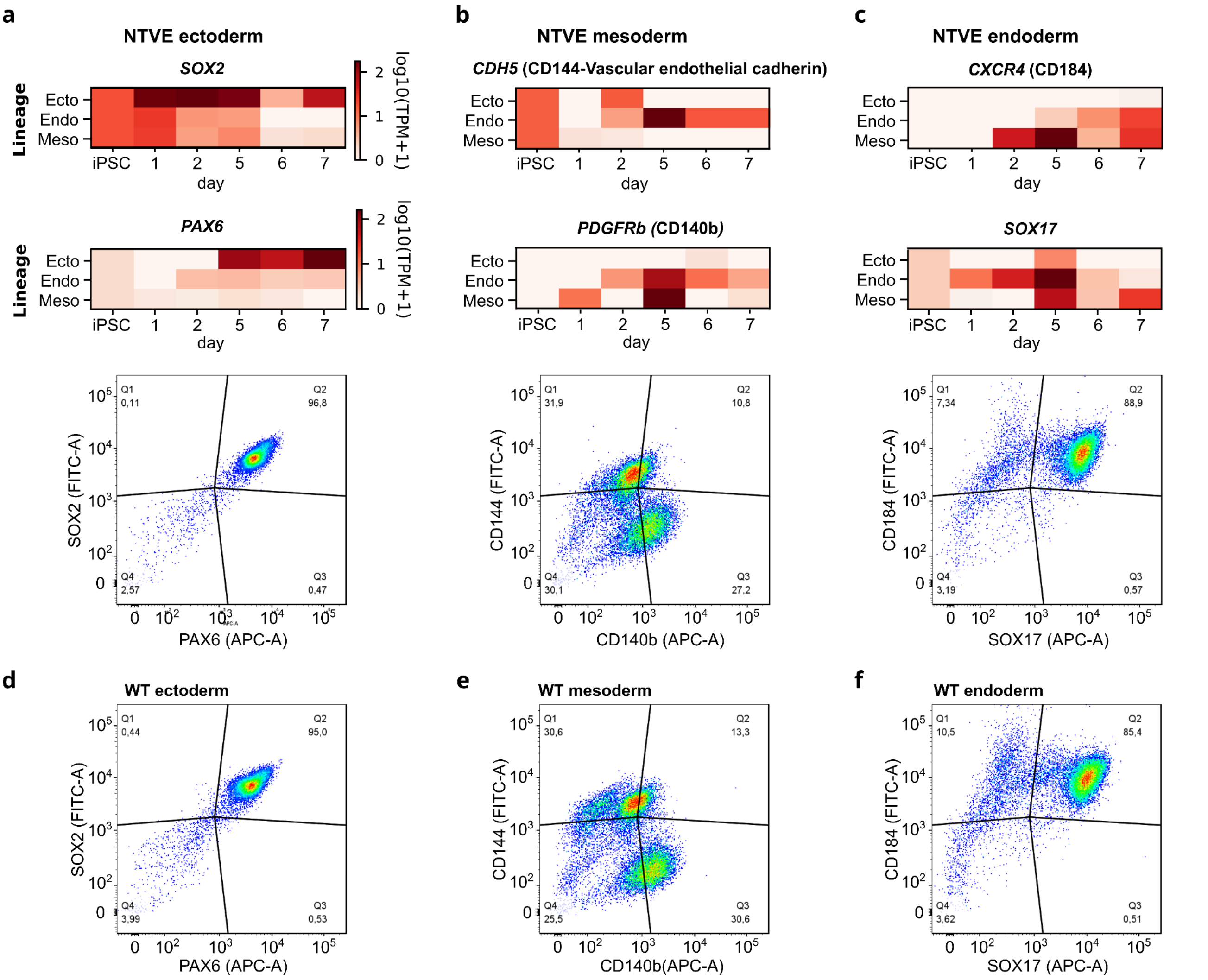


**Supplementary Figure 19: Detection of lineage-specific markers after SCTi003-A hiPSC differentiation.**
**(a,b,c)** NTVE-derived transcript abundance of recommended lineage-specific marker sets for **(a)** ectoderm, **(b)** mesoderm, and **(c)** endoderm (top; *n* = 3), compared with endpoint surface protein detection by flow cytometry using the corresponding antibodies (bottom; *n* = 1). **(d,e,f)** Parental (wild-type) SCTi003-A hiPSCs were analyzed at the end of differentiation (*n* = 1) by immunostaining with lineage-specific markers followed by flow cytometry. Representative gating strategies and expression profiles are shown for **(d)** ectoderm, **(e)** mesoderm, and **(f)** endoderm. Raw sequencing data and analysis code are available *via* Zenodo deposition.

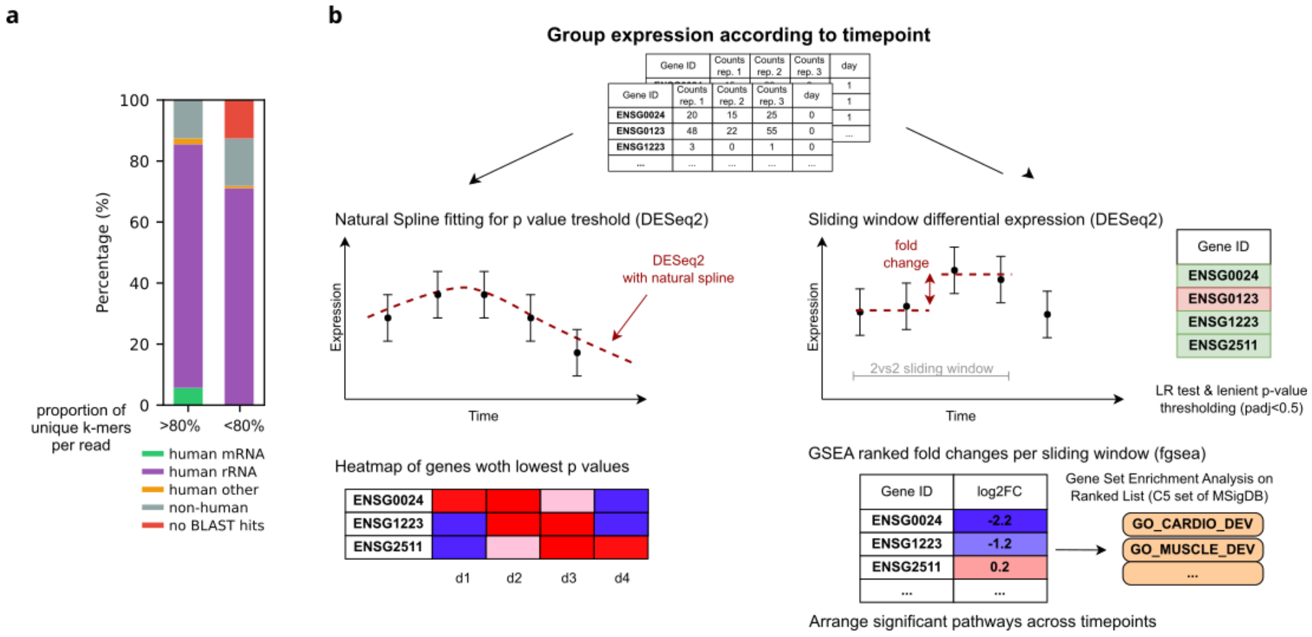


**Supplementary Figure 20: K-mer quality filtering and time-resolved expression analysis of cardiomyocyte differentiation.**
**(a)** Reads from the cardiomyocyte differentiation time course (**Figure 7**) were analyzed using the pipeline described in Supplementary **Figure 10**. To evaluate the effect of the k-mer uniqueness filter, sequences from the FastQC overrepresentation reports were separated into reads that passed (defined as >80% unique k-mers and >80 contiguous bases mapped) and those that were rejected (<80% unique k-mers). For computational tractability, only overrepresented sequences were subjected to BLAST analysis against the NCBI nucleotide database to determine their molecular and taxonomic composition. Results are displayed as stacked bar charts showing the composition of overrepresented sequences, categorized into mutually exclusive groups by BLAST hit identity. Of note, the fraction of human mRNA reads among the overrepresented sequences in the retained set (>80% k-mer uniqueness) is small because most human transcripts are not individually overrepresented; critically, none of these sequences were erroneously assigned to the rejected read set (<80% k-mer uniqueness). **(b)** Time-dependent expression of protein-coding genes was modeled using a natural spline basis, and genes with significantly dynamic expression were identified by a likelihood ratio test (FDR < 0.05). Significant genes were sorted by time to peak expression and visualized as an expression heatmap. Time-resolved differential expression was determined using a sliding window approach, comparing two consecutive days against the subsequent 2 days with a 1-day stride. Raw sequencing data and code are available at Zenodo.

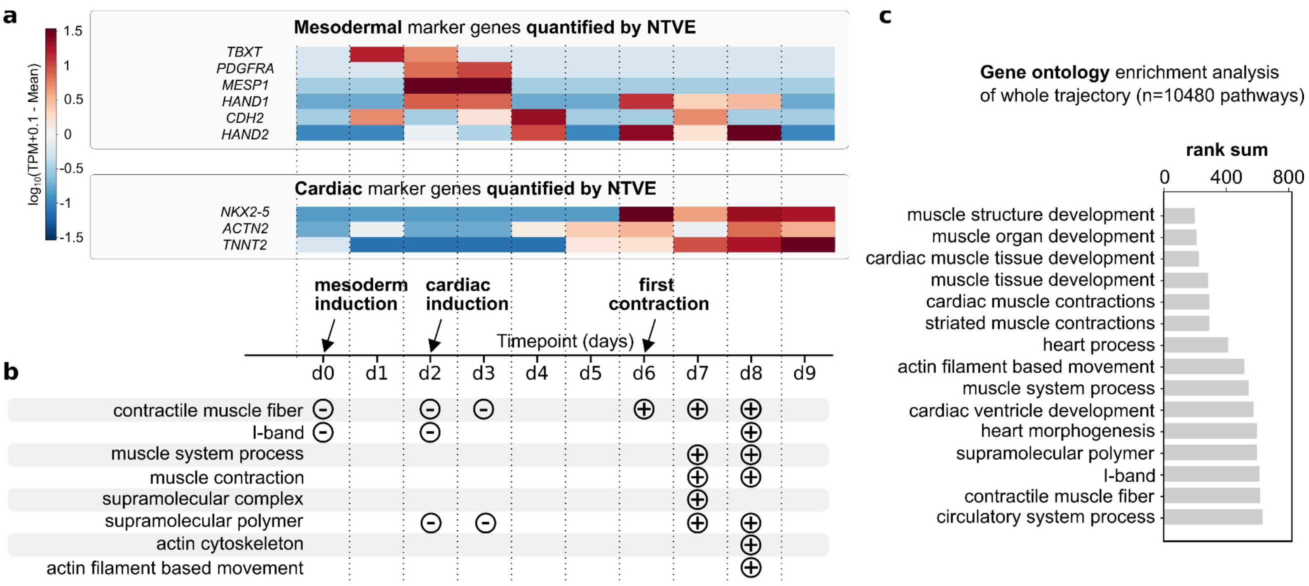


**Supplementary Figure 21: NTVE monitoring of standard marker genes and pathway enrichment over all time points**
**(a)** Mean-centered $\log_{10}$ expression trajectories of established mesodermal and cardiac marker genes detected *via* NTVE over a 9-day differentiation time course in MRIi003-A $NTVE_{PABP}$. **(b)** Day-resolved gene set enrichment analysis (GSEA) of NTVE-derived expression profiles. Expression on each day was compared to the time-course mean; enrichment (+) or depletion (−) of pathways relative to this baseline is indicated. The top significantly enriched or depleted pathways are shown per day. **(c)** For each differentiation day, genes were ranked by expression level and subjected to GSEA against 10,480 curated gene sets. Enrichment scores were compiled across all time points, and pathways were ranked by adjusted *P* value. The top 15 pathways with significant enrichment (FDR < 0.05) across the time course are shown. Raw data and code are deposited at Zenodo.

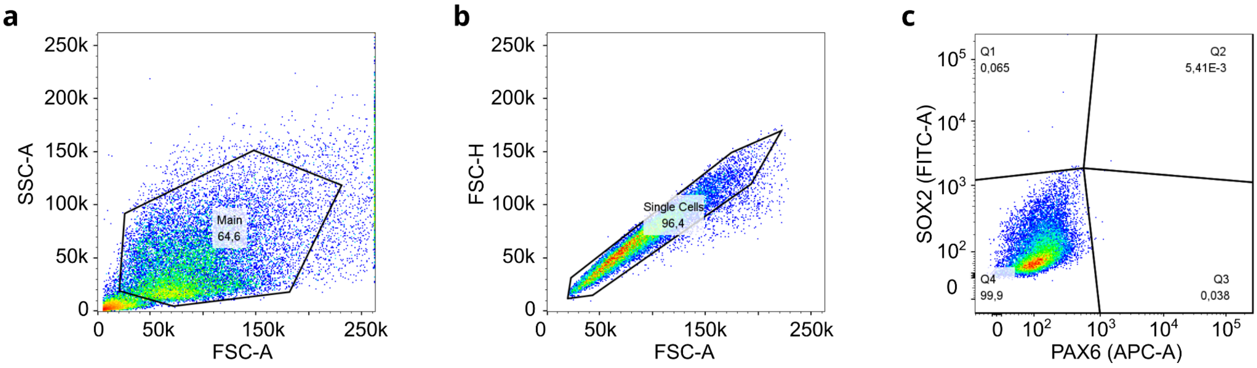


**Supplementary Figure 22: Representative flow cytometry gating strategy.**
**a)** Forward scatter area (FSC-A) versus side scatter area (SSC-A) gating to identify the main cell population. **(b)** FSC-A versus forward scatter height (FSC-H) gating to exclude doublets. **(c)** Fluorescence intensity in the APC-A and FITC-A channels was used to identify fluorescent-positive populations.
